# Leaf age modulates physiological and metabolic responses to contrasting nitrogen forms in Chinese fir (*Cunninghamia lanceolata* (Lamb.) Hook)

**DOI:** 10.64898/2026.02.25.707992

**Authors:** Wenyang Fu, Yali Zhang, Wanting Yu, Zhongwei Zhang, Shu Yuan, Guangdeng Chen, Jian Zeng

**Affiliations:** College of Resources, Sichuan Agricultural University, Wenjiang, 611130, Sichuan, China; International Science and Technology Cooperation Base for Efficient Utilization of Nutrient Resources and Fertilizer Innovation, Sichuan Agricultural University, Wenjiang, 611130, Sichuan, China

**Keywords:** Chinese fir, Leaf age, N forms, Physiological response, Metabolic reprogramming, N addition

## Abstract

Atmospheric nitrogen (N) deposition is increasingly affecting global ecosystems, with nitrate contributing a growing proportion alongside ammonium. However, the interaction between N forms and leaf developmental stage in shaping physiological and metabolic strategies in Chinese fir remains poorly understood. In this study, a field experiment was conducted to explore the physiological and metabolic responses of young and old leaves to ammonium and nitrate N addition. Our findings showed that N addition enhanced photosynthetic performance in young leaves, with a stronger effect from nitrate. In contrast, old leaves exhibited limited photosynthetic response but accumulated higher non-structural carbohydrates and showed elevated N assimilation enzyme activities, particularly under nitrate addition. Phytohormone profiles varied between leaf ages, with young leaves having higher auxin levels while old leaves exhibiting increased abscisic and salicylic acid contents under N addition. Additionally, N addition induced differential reprogramming of amino acid metabolism, with age-dependent accumulation patterns. Metabolomic analysis identified key amino acids involved in coordinating carbon-nitrogen metabolism. These results highlighted the complementary metabolic strategies by young and old leaves of Chinese fir under contrasting N forms addition and emphasized the importance of considering both N form and leaf age in optimizing N management for sustainable plantation practices.

**Highlights:**

1. Nitrate enhanced photosynthesis in young Chinese fir leaves more effectively than ammonium.
2. Old leaves prioritized C storage and N assimilation under N addition, especially nitrate.
3. Complementary metabolic strategies between leaf ages optimized resource use under different N forms addition.

## 1. Introduction

A broad range of global ecosystems have been exposed to atmospheric nitrogen (N) deposition at unprecedented levels, driven predominantly by agricultural practices and industrial processes (Peñuelas et al., 2013). Atmospheric N enters natural and seminatural ecosystem through precipitation and dry deposition, modulating plant growth via nutritional status and photosynthetic processes (Liang et al., 2020; Hu and Liu, 2022; Püspök et al., 2023). In China, atmospheric N deposition has been on the rise since the 1980s and subsequently maintained a high, stable level to date (Yu et al., 2019). N is an indispensable macronutrient for plants, playing a crucial role in the formation of proteins, nucleic acids, chlorophyll, and other bioactive molecules, thereby promoting plant growth and productivity (Raven, 2003). Plants demand N in substantially higher quantities than most other nutrients, making its availability a key factor in primary productivity (LeBauer and Treseder, 2008). In terrestrial ecosystems, plants acquire N in ammonium (NH_4_ ^+^) and nitrate (NO_3_ ^−^) forms, each exerting distinct influences on plant performance due to their divergent metabolic assimilation pathways ( Wang et al., 2018; Kishorekumar et al., 2020). These N forms not only influence plant growth patterns but also impact ecosystem health and stability (Liu et al., 2024). As atmospheric N deposition shifts from ammonium dominance to co-dominance with nitrate (Yu et al., 2019), understanding how different N form affect plant growth and ecosystem stability is essential for optimizing N management strategies in agricultural and ecological systems.

Phytohormones such as auxin (IAA), abscisic acid (ABA), jasmonic acid (JA), salicylic acid (SA), and so on, regulate plant growth and development, mediate responses to environmental changes and influence carbon (C) and N assimilation (Xing et al., 2022; Long et al., 2025). For example, phytohormones can induce the redirection of carbon flux toward the synthesis of carbohydrates and proteins and vary across leaf development stages (Xu et al., 2018; Fàbregas et al., 2026). Furthermore, phytohormones exerts feedback regulation on N metabolism, aligning plant nutrition with growth, development, and growth conditions (Vega et al., 2019).

*Cunninghamia lanceolata* (Lamb.) Hook (Chinese fir) is a fast-growing, ecologically significant timber tree species in China. The issue of productivity decline associated with successive plantings has become increasingly prominent. Within this context, the need for efficient nutrient management strategies is becoming important. N plays a critical role in sustaining plantation productivity, as it affects both soil level and leaf processes. Leaf as the main organ of plant in direct contact with the environment, are predictive of growth performance and ecological function (Falster et al., 2018). Leaf functional traits are sensitive to N availability, especially in ecosystems with low N deposition background (Escudero and Mediavilla, 2003; Knoor et al., 2005). Moreover, the leaf phenology and resource allocation are affected by leaf age, with young leaves undergoing vigorous growth and requiring substantial carbon and N for the biosynthesis of essential compounds ( Yan et al., 2012; Poucet et al., 2021). Old leaves shift toward remobilizing nutrients or resources to support plant growth (Guo et al., 2025). Studies on Arabidopsis and wild soybean highlighted the differential response of young and old leaves to N availability and environmental stress (Berens et al., 2019; Gao et al., 2022). These findings suggested that understanding the age dependent response of young and old leaves to different N forms addition is key to optimizing nutrient management strategies, which is essential for improving the sustainability and productivity of Chinese fir plantations.

Despite the well-established importance of N forms and leaf age, little research has explored how ammonium and nitrate affect the functional traits of young and old leaves in Chinese fir. In this study, a filed-controlled simulation of N addition was performed in Chinese fir plantations with aim of (1) exploring the functional response of young and old leaves to ammonium and nitrate addition, (2) clarifying the regulatory effect of N sources on leaves functional traits and (3) uncovering the specific nutrient strategies during leaf development. These findings will advance the elucidation of adaptive mechanism of Chinese fir to environmental changes, which in turn establishes a robust foundation for developing targeted N management strategies that balance productivity enhancement with ecosystem sustainability.

## 2. Materials and methods

### 2.1. Study region and experiment design

The N addition experiment was conducted at Hongya National Forest Farm in the western Sichuan Province, China (29°38’ N, 102°58’ E), with a humid subtropical climate. The site had an average annual sunlight duration of 1006.1 h, mean annual temperature of 16.6 °C and precipitation of 1453.5 mm. Prior to fertilization addition, the stand was a two-year-old Chinese fir plantation with a density of about 1200 trees per hectare and an average height of around 50 cm.

The experiment used a randomized block design with three blocks, each containing three 15 m × 15 m plots. The basic soil properties were described in a previous study (Yu et al., 2020). The treatments contained nitrate addition (5 g NO_3_^-^ m^-2^ year^-1^), ammonium addition (5 g NH_4_^+^ m^-2^ year^-1^) and a control (no N addition). Treatments were randomly assigned to plots within each block, with a 20 m buffer region between adjacent plots. (NH_4_)_2_SO_4_ and Ca(NO_3_)_2_ was applied to the N addition treatment, and distilled water was used for the control. The treatment lasted from April to October in 2022.

### 2.2. Foliage sampling

At the filed site, Chinese fir leaves were classified into two age groups, current year (young) and two-year-old (old) leaves, distinguished by leaf color and bud scars. The corresponding samples for treatments were as follows, young leaves (YCON) and old leaves (OCON) under control, young leaves (YAN) and old leaves (OAN) under ammonium addition, young leaves (YNN) and old leaves (ONN) under nitrate addition. In October 2022, at least 40 leaves from each age group were randomly sampled from each branch and transported to the laboratory and stored at −80°C for no more than two weeks before physiological, morphological and metabolomic analyses.

### 2.3. Observation of leaf ultrastructure characteristics

Leaves for ultrastructure observation were randomly selected from each treatment and age class. The middle part of leaves was fixed in 3% (v/v) glutaric aldehyde in 0.2 M phosphate buffer (pH 7.2) under negative pressure in a syringe. After fixation, the samples were dehydrated in an ethanol series and embedded in Epoxy Embedding Medium (Sigma-Aldrich Chemie, Steinheim, Germany). Ultrathin sections (80 nm) were cut using a 45° diamond knife on an ultramicrotome and stained with 3% uranyl acetate and 2.7% lead citrate. The sections were then mounted on copper grids and observed under a JEM 2100 plus transmission electron microscopy (JEOL, Tokyo, Japan). Digital images were captured with an EMSIS QUEMESA TEM CCD camera (EMSIS GmbH, Münster, Germany). At least three micrographs per sample were acquired and analyzed to assess ultrastructural changes.

### 2.4. Photosynthetic capacity performance

Due to the small area of individual Chinese fir needle, 4-5 needles of the same age class were randomly selected and placed in the gas exchange chamber. Leaf photosynthesis rate was measured using a LI-6400 portable infrared gas analyzer (LI-Cor Biosciences, Lincoln, NE, USA) to produce light response curves in descending order of 2000, 1800, 1600, 1400, 1200, 1000, 800, 600, 400, 200, 100, 50 and 0 μmol m^-2^ s^−1^. Prior to measurements, leave were acclimated in the chamber at 25°C, with ambient humidity and 400 μmol mol^−1^ CO_2_ for 30 min to stabilize leaf photosynthesis. The net photosynthetic rate (*P*_n_) was automatically recorded at 3-min intervals at each light level. Maximum net photosynthesis (*P*_max_) was determined from the plateau in the light response curves and the rectangular hyperbola model for *P*_max_, dark respiration rate (*R*_d_), light compensation point (*L*_CP_), light saturation point (*L*_SP_) and apparent quantum yield (AQY) (Kyei-Boahen et al., 2003).

### 2.5. Analysis of leaf physiological functional traits

Non-structural carbohydrates (NSC) were determined by the phenol-sulfuric acid method (Buysse and Merckx, 1993). Plant samples was extracted with 80% alcohol to obtain the supernatant for soluble sugar determination, followed by extraction of residues with 1 M H_2_SO_4_ to determine starch after a water bath and centrifugation. Soluble sugar and starch were measured spectrophotometrically at 625 nm using the enthrone reagent. NSC was calculated as the sum of soluble sugar and starch concentrations. Rubisco activity was determined by monitoring NADPH oxidation at 340 nm (Sales et al., 2020). Nitrate reductase (NR) activity and nitrite reductase (NiR) activity were determined using commercial assay kits (ARG82016 and ARG83413, Arigo Biolaboratories Corp., Shanghai, China). Glutamine synthetase (GS) and Glutamate synthase (GOGAT) activities were measured using commercial kits (Beijing Solarbio Science & Technology Co., Ltd., Beijing, China) via spectrophotometry (GS: Cat. BC0915; GOGAT: Cat. BC0075). Total free amino acids were extracted by ultrasonication with 5% (v/v) aqueous trifluoroacetic acid followed by ultrafiltration and detection by high performance liquid chromatography (Zheng et al., 2017).

### 2.6. Determination of phytohormone contents

Phytohormone contents in leaf samples were determined using established methods (Pan et al., 2010). IAA content was measured by the phenol–sulfuric acid method after extraction with 95% ethanol and quantified by spectrophotometric absorbance using a standard curve. JA content was determined by gas chromatography after extraction with acetonitrile: water (1:1, v/v) and ultrasonic treatment. ABA content was quantified using an enzyme-linked immunosorbent assay (ELISA) according to the manufacturer’s instructions. SA content was analyzed by high-performance liquid chromatography following liquid N grinding and extraction.

### 2.7. Metabolome profiling

Freeze-dried biological samples were ground into fine powder using a mixer mill (MM 400, Retsch) at 30 Hz for 1.5 min. Approximately 100 mg of the powdered sample was dissolved in 1.0 mL of extraction solution (70% methanol containing 0.1 mg L^-1^ lidocaine as an internal standard). The mixture was incubated overnight at 4 °C, vortexed three times to improve extraction efficiency. After centrifugation at 10,000 g for 10 min, the supernatant was collected, filtered through a 0.22 μm microporous membrane, transferred into sample vials, and stored for analysis.

Metabolite detection was performed using UPLC-MS/MS (UPLC–MS/MS; Thermo Fisher Scientific, Waltham, MA, USA), and the quantification, and identification were conducted by Wuhan Metware Biotechnology Co., Ltd. Metabolite identification was based on public databases, including MassBank, LipidMaps, mzCloud, and Kyoto Encyclopedia of Genes and Genomes (KEGG) with supplementary confirmation using Human Metabolome Database (HMDB). Principal component analysis (PCA) and partial least squares discriminant analysis (PLS-DA) were applied to evaluate metabolite accumulation patterns using the R package ‘ropls’ (version 4.3.1). Differentially accumulated metabolites (DAMs) were identified based on variable importance in projection (VIP) values > 1 and |log_2_ fold change| > 1. K-means clustering analysis in R package ‘stats’ (version 4.3.1) was grouped DAMs into subclasses, with the optimal clusters number determined by the Calinski-Harabasz index. KEGG pathway enrichment was conducted for DAMs in each subclass to identify significantly enriched metabolic pathways. Metabolic pathway and pathway-metabolite networks were constructed using Cytoscape software (3.10.3). Network topology analysis was performed to assess pathway-metabolic interactions.

### 2.8. Statistical analysis

One way analysis of variance in SPSS 27.0 was used to analyze the significance of differences (*p* < 0.05) between leaf ages in the physiological data across contrasting N forms addition. Graphs were plotted using Origin 2018 software.

## 3. Results

### 3.1 Photosynthetic performance and C metabolism under contrasting N forms addition

Distinct differences in chloroplast ultrastructure were observed between young and old leaves of Chinese fir under different N sources addition (Fig. 1). In young leaves, chloroplasts were predominantly spindle-shaped and closely associated with the cell wall. Thylakoid structure was orderly arranged, and the stromal lamellae were tightly packed and clearly visible. NO□□–N promoted greater starch accumulation than NH□□–N (Fig. 1A-C). In old leaves, chloroplast ultrastructure varied among N treatments, characterized by cell wall thickening, granum disintegration, peripheral reticulum roughening, and starch accumulation, with NH□□–N notably inducing starch granule dilation (Fig. 1D–F).

**Figure 1.**
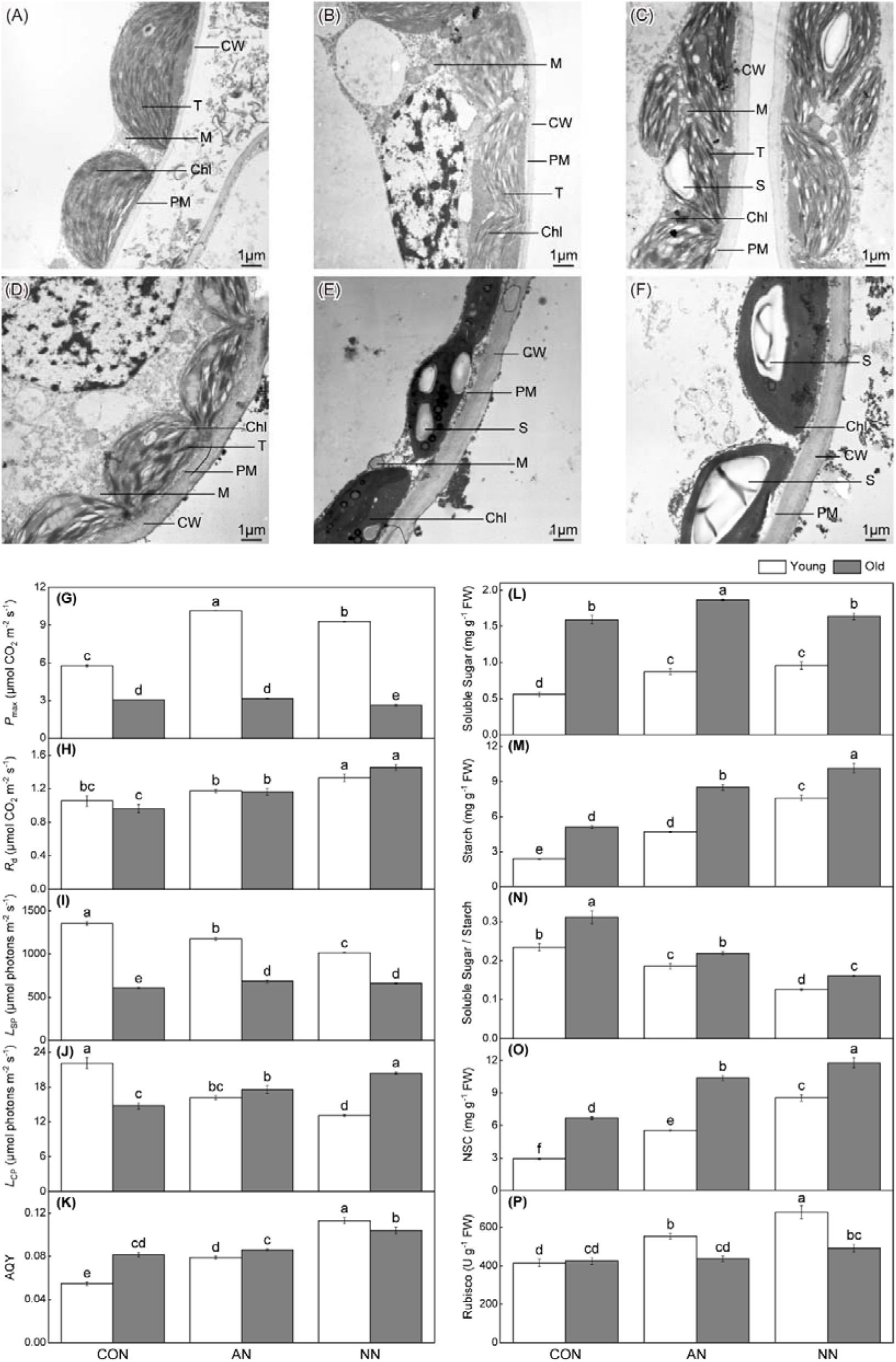
Differential effects of contrasting N forms addition on photosynthetic potential in young and old leaves of Chinese fir. Transmission electron microscopy images of chloroplast ultrastructure in young leaves (A–C) and old leaves (D–F) under control without N addition (CON), NH_4_^+^–N addition (AN) and NO_3_^-^–N addition (NN). Photosynthetic parameters and carbohydrate-related traits in young (white bars) and old leaves (grey bars), including *P*_max_, maximum net photosynthetic rate (G); *R*_d_, Dark respiration rate (H); LSP, Light saturation point (I); LCP, Light compensation point (J); AQY, Apparent quantum yield (K); Soluble sugar content (L); Starch content (M), Soluble sugar/starch ratio (N); Non-structural carbohydrate content (O); Rubisco activity (P). Different lowercase letters indicated significant differences among treatments and between leaf ages (*p* < 0.05). Scale bars = 1 μm. CW, cell wall; PM, plasma membrane; Chl, chloroplast; M, mitochondria; T, thylakoid; S, starch granule.

Photosynthetic performance and associated C metabolism also differed markedly between young and old leaves under contrasting N forms addition (Fig. 1G–K). *P*_max_ and *L*_SP_ were consistently higher in young leaves. N addition enhanced Pmax in young but reduced it in old leaves. *R*_d_ and AQY increased in both leaf types following N addition. *L*_CP_ and *L*_SP_ decreased in young leaves but increased in old leaves under N addition, although *L*_SP_ remained higher in young leaves overall.

There were significant differences in the carbohydrate accumulation and allocation patterns between young and old leaves under N additions (Fig. 1L-P). Old leaves contained higher soluble sugar, starch, and NSC contents. N addition increased carbohydrate levels in both ages but reduced the soluble sugar/starch ratio. NO□□–N resulted in greater carbohydrate accumulation than NH□□–N, whereas the sugar/starch ratio showed the opposite trend. Rubisco activity increased significantly in young leaves under N addition, particularly with NO□□–N, but showed little change in old leaves.

### 3.2 N metabolism under contrasting N forms addition

N metabolism-related enzymes activities and free amino acid content showed an increasing trend under N addition except for GS activity in young leaves (Fig. 2). Enzyme activities, including NR, NiR, GOGAT, GS and GDH, were significantly higher in old leaves than in young leaves. Correspondingly, free amino acid content was also higher in old leaves, irrespective of N forms. Compared to NH_4_^+^–N addition, NO_3_^-^–N addition significantly enhanced NR and NiR activities (Fig. 2A, B). The activities of GS, GOGAT and GDH varied differently between young and old leaves under N addition. However, there were no significant difference in free amino acid content between N forms for both leaf ages (Fig. 2F).

**Figure 2.**
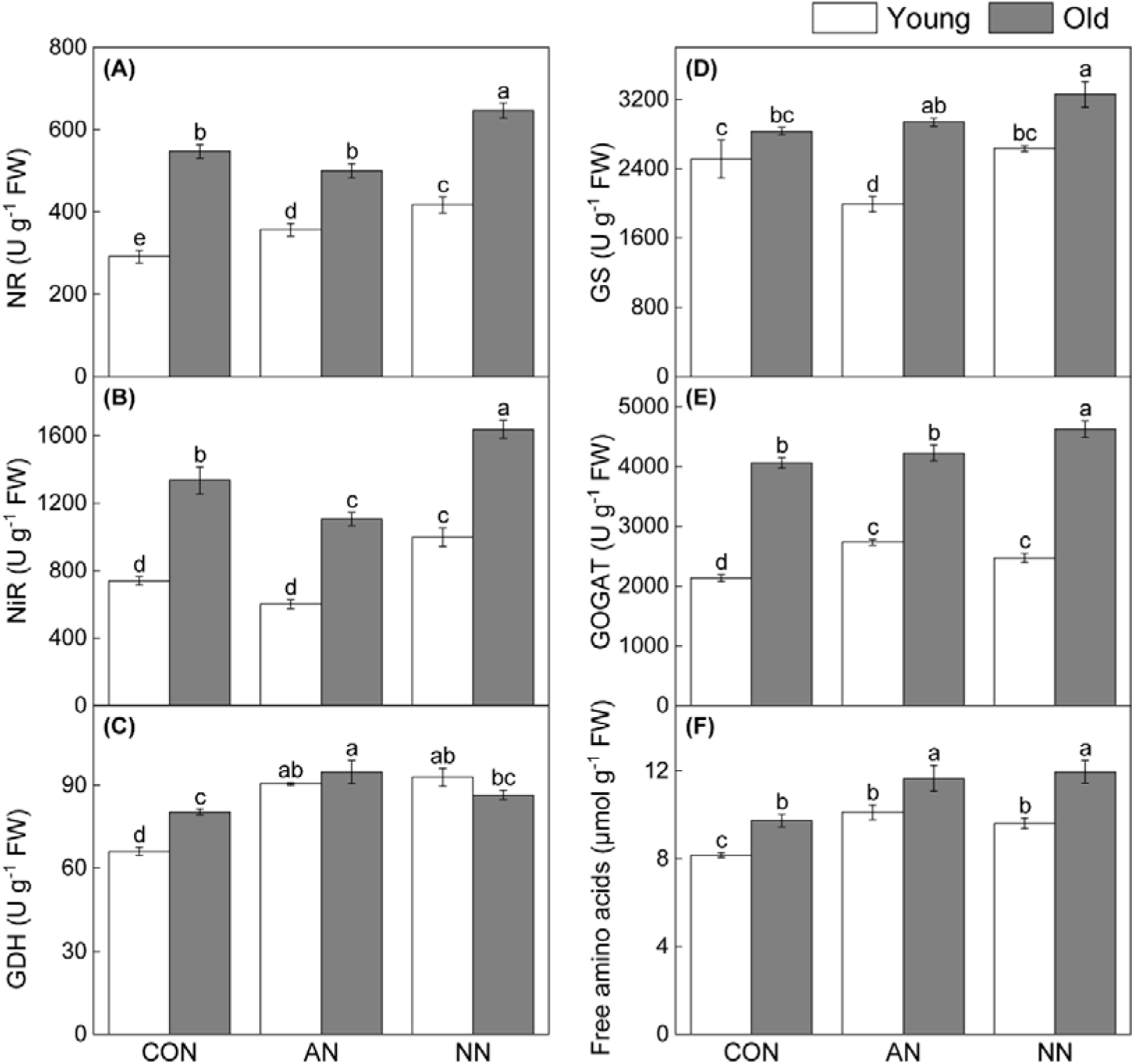
Effects of contrasting N forms addition on N metabolism in young and old leaves of Chinese fir. Nitrate reductase activity (A), Nitrite reductase activity (B), Glutamate dehydrogenase activity (C), Glutamine synthetase activity (D), Glutamate synthase activity (E), Free amino acid content (F). Different lowercase letters indicated significant differences among treatments and between leaf ages (*p* < 0.05).

### 3.3 Phytohormone profiling under contrasting N forms addition

Phytohormone contents varied significantly between young and old leaves in response to different N forms. Except for IAA, JA, ABA and SA contents were consistently higher in old leaves than in young leaves (Fig. 3). In young leaves, N addition significantly increased IAA and JA but decreased ABA and SA compared to control. In old leaves, NH_4_^+^–N addition significantly decreased IAA, JA and SA contents but increased ABA level, whereas NO_3_^-^–N addition significantly elevated IAA, ABA and SA levels compared to control (Fig. 3A, C, D). Compared with NH□□–N, NO□□–N significantly decreased JA and ABA but increased SA in both ages. IAA content was significantly enhanced by NO_3_^-^–N addition only in old leaves (Fig. 3). Notably, NO_3_^-^–N strongly enhanced age-dependent differences in ABA and SA accumulation, whereas NH_4_^+^–N exerted a weaker inductive effect on these hormones in old leaves.

**Figure 3.**
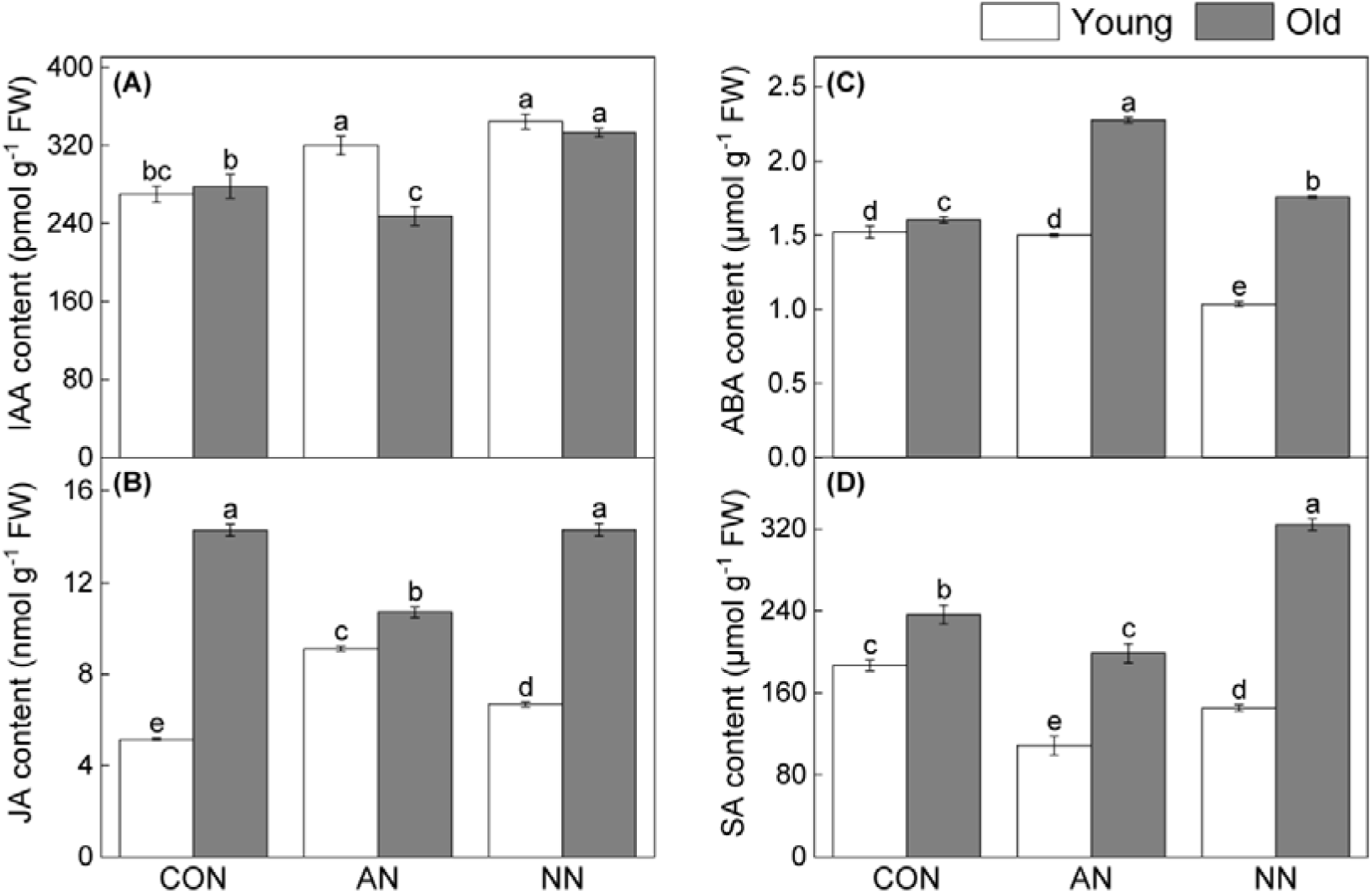
Effects of contrasting N forms addition on phytohormone contents in young and old leaves of Chinese fir. Auxin content (A), Jasmonic acid content (B), Abscisic acid content (C), Salicylic acid content (D). Different lowercase letters indicated significant differences among treatments and between leaf ages (*p* < 0.05).

### 3.4 Metabolites accumulation patterns under contrasting N forms addition

To characterize metabolic pathway responses to contrasting N forms and leaf ages, metabolomic profiling was conducted (Fig. 4). PCA and PLS-DA analyses revealed clear separation according to leaf age and N form (Fig. 4A, B). A total of 562 metabolites were detected. Compared with control, there were 53 DAMs (36 up-regulated, 17 down-regulated) in YAN, 43 DAMs (34 up-regulated, 9 down-regulated) in YNN, 40 DAMs (20 up-regulated, 20 down-regulated) in OAN, and 32 DAMs (26 up-regulated, 6 down-regulated) in ONN (Fig. 1C). DAMs were primarily enriched in pathways related to C and N metabolism, including amino acidmetabolism, translation, membrane transport, and secondary metabolites biosynthesis (Fig. S1). Age-dependent enrichment patterns were observed (Fig. S1A–D). Young leaves showed enrichment primarily in amino acid biosynthesis and carbon metabolism, especially aminoacyl-tRNA and glucosinolate pathways (Fig. S1A, B) In old leaves, NH_4_ ^+^–N addition resulted in comparatively limited enrichment, largely confined to amino acid and nucleotide metabolism (Fig. S1C). In contrast, NO_3_^-^–N addition induced broader metabolic adjustments, particularly in secondary metabolism, with higher enrichment in anthocyanin biosynthesis (Fig. S1D). NH_4_^+^–N specifically enriched ABC transporters and monobactam biosynthesis (Fig. S1E), whereas NO_3_^-^–N uniquely enriched carbapenem biosynthesis and 2-Oxocarboxylic acid metabolism in old leaves (Fig. S1F). Furthermore, N addition generated 15 unique DAMs in young leaves and 22 in old leaves compared to control (Fig. 1D, E). Unique DAMs in young leaves were mainly flavonoids, organic acids, and plant hormones, while old leaves exhibited a more diverse DAM profile, additionally including amino acids. Relative to young leaves, old leaves exhibited 17 unique DAMs under N addition. Under NH_4_^+^–N addition, old leaves showed enrichment in fatty acids, amino acids, and flavonoids, while NO_3_^-^–N addition uniquely accumulated nucleotides and their derivatives (Table S1, S2).

**Figure 4.**
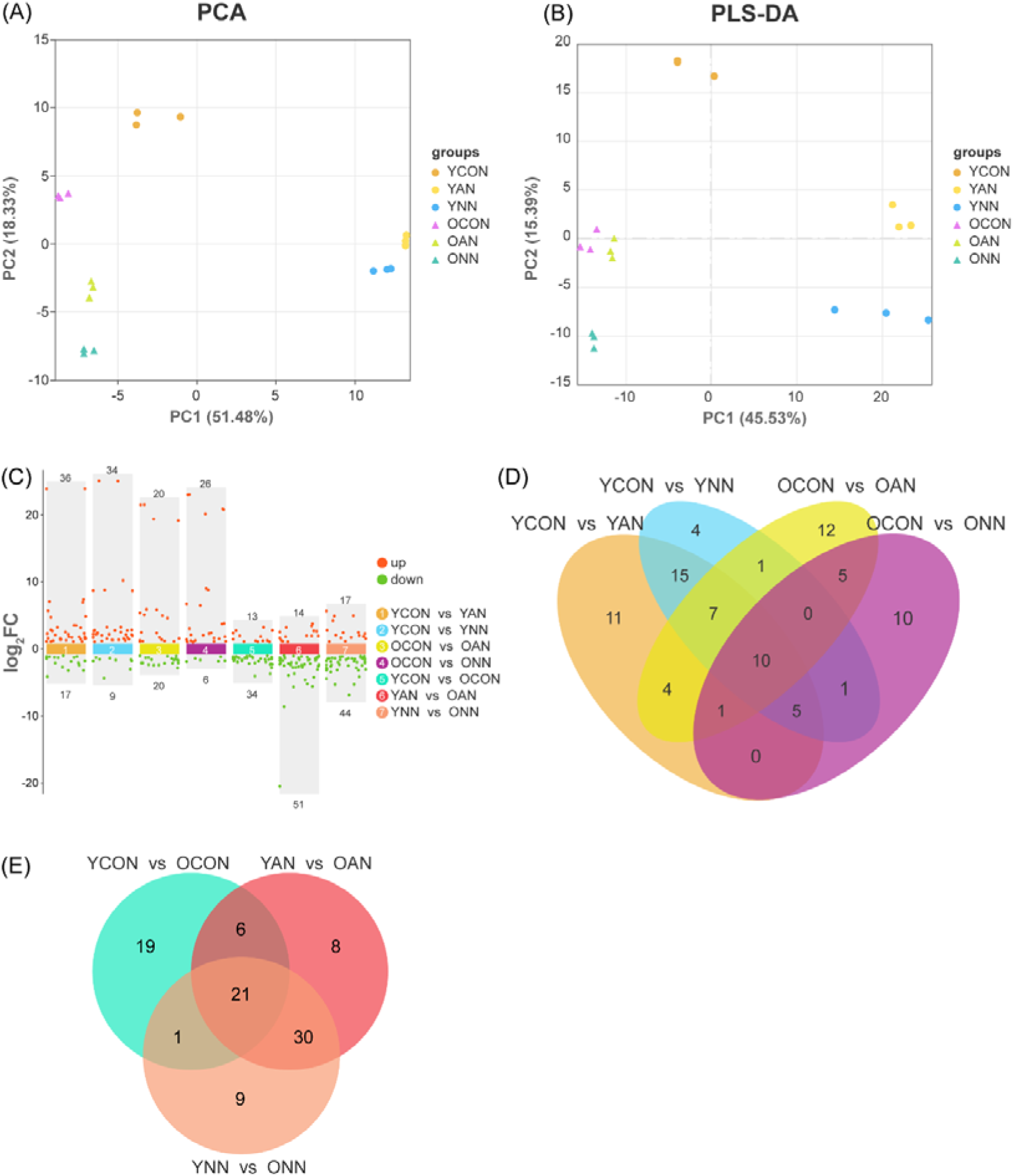
Multivariate analysis of metabolomics data and identification of differentially accumulated metabolites (DAMs). Principal component analysis (PCA) score plot of metabolome data (A), Partial least squares discriminant analysis (PLS-DA) score plot (B), Distribution of DAMs identified in pairwise comparisons (C), Venn diagrams illustrated the shared and unique DAMs under differential N forms addition (D). Venn diagrams illustrated the shared and unique DAMs between different leaf types (E). YCON, young leaves under control conditions without N addition; YAN, young leaves under NH_4_^+^–N addition; YNN, young leaves under NO_3_^-^–N addition; OCON, old leaves under control conditions without N addition; OAN, old leaves under NH_4_^+^–N addition; ONN, old leaves under NO ^-^–N addition.

### 3.5 DAMs clustering and metabolic pathway response to contrasting N form addition

K-means clustering identified five distinct DAMs clusters across N forms treatments and leaf ages (Fig. 5A–F). Cluster 1 was specifically upregulated in young leaves under NO_3_^-^–N addition, while Cluster 3 increased in N-treated young leaves but decreased in old leaves. Other clusters displayed treatment- or age-specific patterns. KEGG enrichment analysis revealed functional differentiation among clusters (Fig. 5G–J). Clusters 1 and 3 were enriched in amino acid-related pathways, including amino acids biosynthesis, aminoacyl-tRNA biosynthesis, and 2-oxocarboxylic acid metabolism. Cluster 1 was additionally enriched in ABC transporters and glucosinolate biosynthesis, while Cluster 3 was characterized by cofactors biosynthesis. Cluster 2 and Cluster 5 were specifically enriched in anthocyanin biosynthesis and purine metabolism, respectively, with no significant pathways detected in Cluster 4. Network analysis highlighted aminoacyl-tRNA biosynthesis and ABC transporters as hub pathways in Cluster 1, with L-arginine, L-tyrosine, L-ornithine, and L-isoleucine exhibiting high connectivity (Fig. 5K). In Cluster 3, cofactor biosynthesis emerged as the central hub, with L-glutamate and L-valine as key nodes (Fig. 5L). Based on network topology and biological relevance, these metabolites were selected as primary candidates. Representative metabolites from Cluser 2 and 5, including pelargonin, keracyanin, 3′-adenylic acid, adenylic acid, guanine, and chlorogenic acid, were also retained. Twelve key metabolites were integrated into a heatmap and correlated with physiological traits (Fig. 5M), revealing significant associations with multiple C- and N-related parameters, thereby linking metabolic shifts to C–N regulation under contrasting N forms addition.

**Figure 5.**
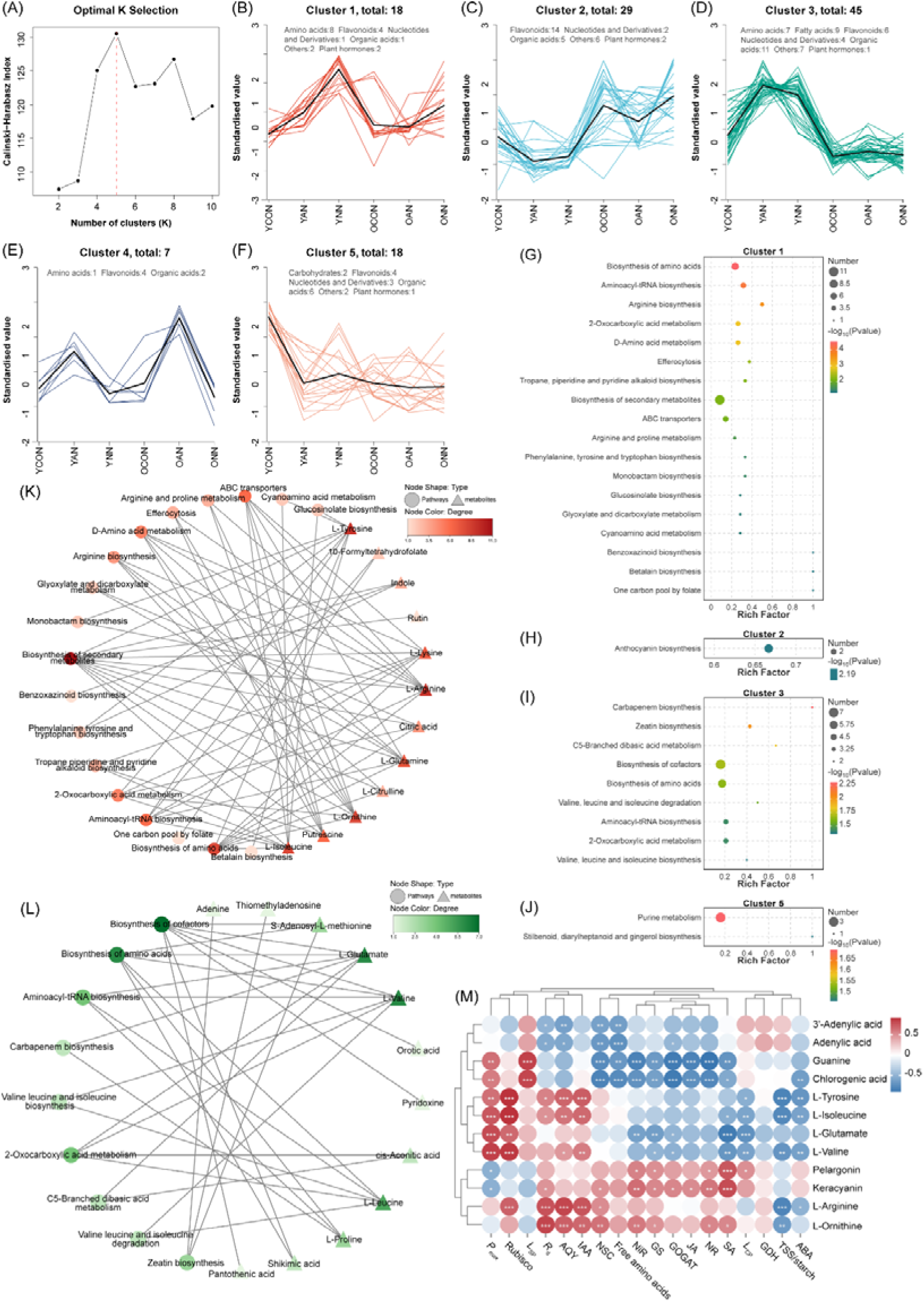
Metabolic clustering patterns and functional associations of differential accumulated metabolites (DAMs) in young and old leaves of Chinese fir. Determination of the optimal number of clusters for K-means clustering (A). K-means clustering profiles of DAMs across treatment and leave ages (B-F). Colored lines represented individual metabolites, and black lines indicated the average trend for each subclass. KEGG pathway enrichment analysis of DAMs in K-means clusters 1, 2, 3, and 5. The rich factor represented the ratio of the number of metabolites mapped to a given pathway to the total number of annotated metabolites (G–J). Bubble size indicated the number of metabolites enriched in each pathway, and bubble color reflected the enrichment significance level. Pathway–metabolite network analyses of enriched metabolic pathways in K-means cluster 1 (K) and cluster 3 (L). Circular and triangular nodes represented metabolic pathways and metabolites, respectively. Node color intensity corresponded to degree values, with darker colors indicating higher connectivity. Pearson correlation analysis between key DAMs and physiological indicators (M). Circle color represented the strength and direction of correlations (red, positive; blue, negative), and asterisks denoted statistical significance (*, *p* < 0.05; **, *p* < 0.01; ***, *p* < 0.001). YCON, young leaves under control conditions without N addition; YAN, young leaves under NH_4_^+^–N addition; YNN, young leaves under NO_3_^-^–N addition; OCON, old leaves under control conditions without N addition; OAN, old leaves under NH_4_^+^–N addition; ONN, old leaves under NO ^-^–N addition.

Mapping these metabolites onto a reconstructed pathway network further clarified age-dependent metabolic reprogramming (Fig. 6). In young leaves, aromatic amino acids (L-tyrosine, L-valine and L-isoleucine) were preferentially upregulated, with stronger accumulation under NO_3_^-^–N addition than NH_4_ ^+^–N addition. In old leaves, amino acids associated with N allocation (L-glutamate, L-arginine and L-ornithine) contents were enhanced, aligning with elevated N assimilation enzyme activities, particularly under NO_3_^-^–N addition. Conversely, purine metabolism exhibited a reduction in nucleotide-related metabolites (3′-adenylic acid, adenylic acid, guanine) under N addition, especially in old leaves. Phenylpropanoid and anthocyanin metabolites were more substantially reconfigured in old leaves, with NO□□–N having stronger a effect on anthocyanin-related compounds.

**Figure 6.**
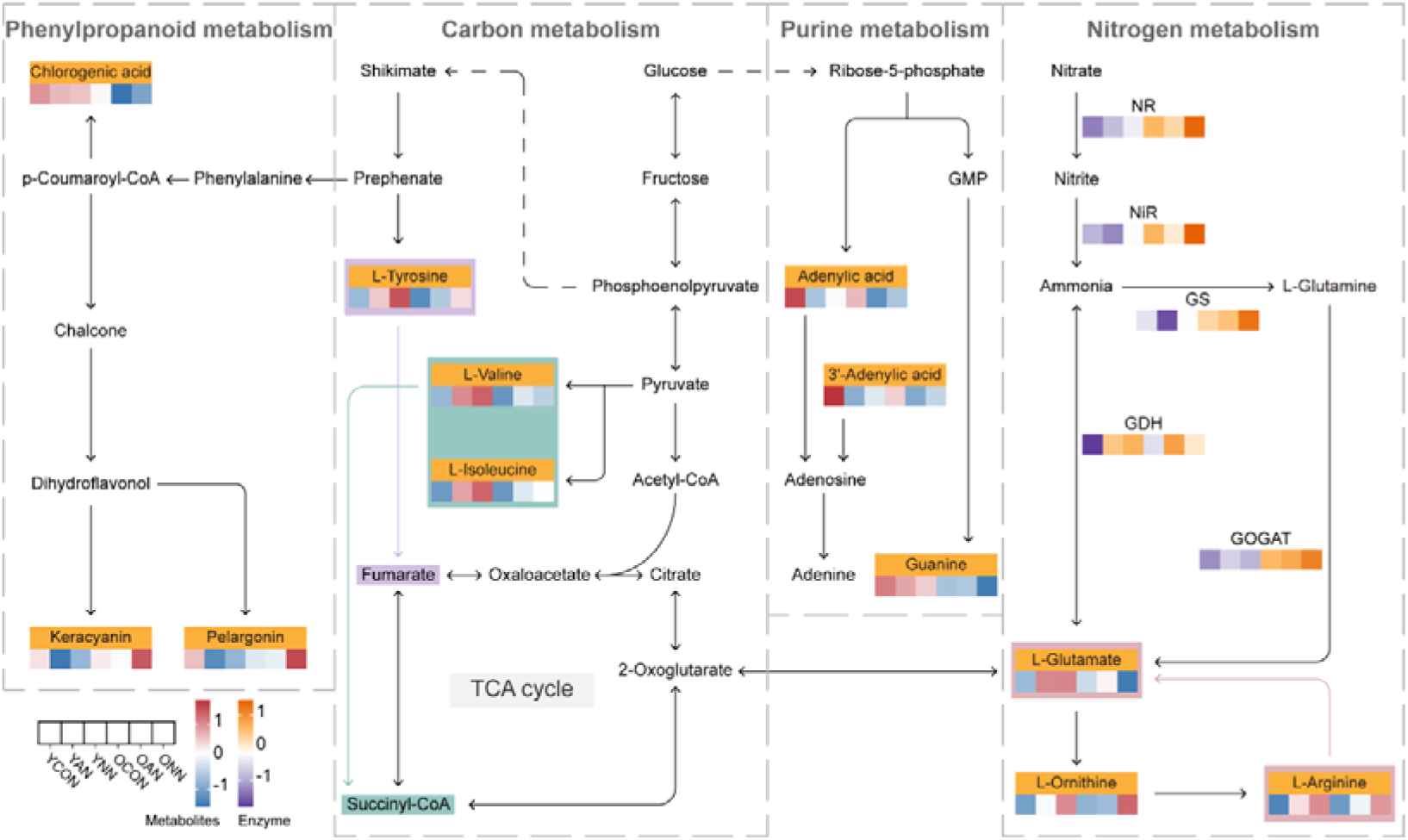
Schematic diagrams of key metabolites involved in N assimilation and amino acid biosynthesis in young and old leaves of Chinese fir under contrasting N forms addition. Heat maps were generated based on row-wise z-score standardized values. Red or blue indicates the mean metabolite abundance, while orange or purple represents the enzyme activity. YCON, young leaves under control conditions without N addition; YAN, young leaves under NH_4_^+^–N addition; YNN, young leaves under NO_3_^-^–N addition; OCON, old leaves under control conditions without N addition; OAN, old leaves under NH_4_^+^–N addition; ONN, old leaves under NO_3_^-^–N addition.

## 4. Discussion

Leaf functional traits varied with different ages or positions, affecting substances and energy exchange between plants and environments, as well as plant survival strategies under changing environmental conditions (Menezes et al., 2022). Sensing nutritional status and signal transduction within the cells and between organs, is essential for coordinating metabolic pathways that dynamically regulate C and N partitioning and utilization (Artins et al., 2024). This study integrates physiological and metabolomic data in Chinese fir to characterize leaf age-dependent response to different N forms addition.

### 4.1 C and N metabolism variation in young and old leaves

Plant organs dynamically adjusted their morphology and function, which in turn affected their nutritional demands (He et al., 2025). C and N are fundamental to plant metabolism, growth and acclimation to environment changes (Nunes-Nesi et al., 2010; Oldroyd and Leyser, 2020). Many key ecological processes varied with leaf age, which is crucial for interpreting plant developmental based on age-dependent leaf traits (Wu et al., 2016). In this study, we demonstrated that leaf age driven profound divergence in photosynthetic C metabolism, N assimilation, phytohormone profile, and metabolite reprogramming under ammonium and nitrate addition, with nitrate generally exerting stronger regulatory effects than ammonium. These findings highlighted an age□dependent C–N coupling mechanism that mediated adaptive responses to contrasting N forms addition in Chinese fir. Young leaves maintained intact chloroplast ultrastructure, higher photosynthetic capacity, and enhanced Rubisco activity following N addition, supporting their role as the primary site for C fixation and growth. By contrast, old leaves exhibited deteriorated thylakoid structure, reduced photosynthetic plasticity, and preferential accumulation of soluble sugars, starch, and NSC, indicating a functional shift toward C storage rather than assimilation. Previous studies revealed that leaf age influenced photosynthetic capacity and feedback inhibition caused by sink limitation was also responsible for age-associated decline in photosynthesis (Han et al., 2008; Tejera et al., 2022). Consistent with structural and physiological observations, nitrate was more effective than ammonium in promoting photosynthetic performance and carbohydrate accumulation, particularly in young leaves. These changes enhanced the adaption ability of young leaves to low light and reduced photoinhibition risk under N addition.

N addition also enhanced light reaction efficiency and intracellular substance synthesis for promoting plant growth and competitive advantage (Shen et al., 2024; Zuo et al., 2026). Our results expanded the understanding of stronger photosynthetic capacity and C gain in young leaves, especially under nitrate addition. Similar contrasts in respiratory costs and photosynthetic plasticity have been described in conifer foliage, where older leaves showed higher maintenance respiration and constrained light-use adjustment (Jensen et al., 2015). This divergence suggested that nitrate facilitated more efficient C capture and provides sufficient C skeletons for N assimilation.

Meanwhile, N addition enhanced the primary C backbone for N uptake, which was evidence by increased N metabolism in both young and old leaves (Fig. 2). However, N metabolism was highly age□dependent, with old leaves displaying significantly higher activities of N assimilating enzymes and greater free amino acid accumulation than young leaves. These results indicated that old leaves were crucial for N assimilation and internal recycling. Compared to ammonium, nitrate had a stronger stimulating effect on NR and NiR activities, reinforcing its role in promoting amino acid metabolism. This supported the view that nutrients were absorbed in younger organs and stored or transported in older organs (Y. Wang et al., 2024). Phytohormones, which modulated C allocation and N uptake, were crucial for plant growth regulation and stress responses (Jia et al., 2022). In this study, N addition increased IAA levels and decreased SA levels, correlating with enhanced photosynthetic capacity and free amino acid accumulation in young leaves (Fig. 3). Ammonium supply not only promoted lateral root emergence by influencing shoot-derived auxin mobility, but also stimulated ABA accumulation for N assimilation and stress adaption (Meier et al., 2020; Sun et al., 2020). In old leaves, higher concentrations of JAs, ABA and SA were accumulated, with nitrate strongly amplifying age□dependent differences in ABA and SA accumulation but ammonium exerting a weaker effect (Fig. 3). This is consistent with previous report that nitrate addition significantly increased SA content in old leaves, which was associated with optimizing C-N metabolism and supporting auxin-mediated growth processes (Xing et al., 2022).

### 4.2 N form addition shapes C-N metabolism coordination between young and old leaves

N availability directly modulates photosynthetic efficiency and C fixation, whereas C status in turn affects N uptake and assimilation. Maintaining a balance between C and N is critical for optimal growth and productivity in plants (Liu et al., 2025). Y. Wang et al. (2024) found that young leaves employed a nutrient acquisition strategy while old leaves focus on nutrient storage strategy. The physiological alternation observed in N addition treated young and old leaves were driven by coordinated biochemical and metabolic changes in C and N metabolism. Young leaves exhibited enhanced photosynthetic performance and amino acid accumulation, whereas NSC storage remained comparatively lower than old leaves (Fig. 1). NSC, consisting of soluble sugar and starch, generally functions as a source of stored energy and C source for biosynthesis (Dietze et al., 2014; MacNeill et al., 2017). The soluble sugar/starch ratio can reflect NSC allocation pattern in leaves, with starch functioning as a transient reserve to balance source activity and sink demand (Martínez-Vilalta et al., 2016; Smith and Zeeman, 2020). Old leaves consistently maintained higher levels of soluble sugars, starch and NSC, alongside stronger N assimilation, indicating its role in coordinating C storage and N assimilation to stabilize metabolic homeostasis and sustain whole plant growth under variable N conditions.

Plants displayed divergent growth responses to different N forms and nitrate or ammonium differentially regulated plant growth and metabolic processes (Bai et al., 2021). Metabolomic analysis revealed age□dependent metabolic reprogramming under contrasting N forms addition. Young leaves preferentially enhanced aromatic and branched□chain amino acids biosynthesis under nitrate addition, supporting growth□related metabolism. This was consistent with that aromatic amino acid biosynthesis is tightly regulated to balance C investment required for rapid growth and downstream biosynthesis (EI-Azaz and Maeda, 2025). In contrast, old leaves exhibited prominent upregulation of N□storage amino acids and secondary metabolites, particularly anthocyanins, reflecting a shift toward stress adaptation and resource conservation. This pattern was more indicative of nitrate-specific metabolic reprogramming rather than N deficiency (Ruffel et al., 2025). Enhanced N assimilation induced by nitrate addition increased the demand for reducing power, stimulating phenylpropanoid and anthocyanin biosynthesis as protective metabolic sinks. Additionally, arginine, ornithine and glutamate were prominent in old leaves, supporting their role in N storage and transport (Ávila et al., 2025). Network analysis identified aminoacyl□tRNA biosynthesis, ABC transporters, and cofactor biosynthesis as hub pathways, with several amino acids and nucleotides serving as key candidates linking metabolic regulation to C–N physiological traits. Purine metabolism was coordinately down□regulated under N addition, especially in old leaves, implying accelerated nucleotide turnover and enhanced N reallocation.

In conclusion, our results reveal a coordinated leaf age-dependent regulatory framework in response to contrasting N forms. Young leaves prioritized photosynthetic C fixation, growth-related amino acid metabolism, and IAA-mediated growth processes, whereas old leaves were characterized by enhanced N assimilation, C storage, and adaptation responses. Nitrate triggers more extensive physiological and metabolic reprogramming than ammonium, thereby strengthening the functional divergence between young and old leaves. This study provided insights into nutrient strategies in leaves of woody plants under heterogeneous N environments.

## Authorship contribution statement

Wenyang Fu: Methodology, Visualization, Validation, Writing–original draft. Yali Zhang: Software, Formal analysis. Wanting Yu: Visualization, Validation. Zhongwei Zhang: Formal analysis, Data curation. Shu Yuan: Writing–review & editing. Guangdeng Chen: Writing–review & editing, Funding acquisition. Jian Zeng: Supervision, Funding acquisition, Writing–review & editing.

## Data Availability Statement

The metabolic datasets generated and analyzed during this study are available in the OMIX database under accession number OMIX015273.

